# Lineage-tracing reveals an expanded population of NPY neurons in the inferior colliculus

**DOI:** 10.1101/2024.03.27.587042

**Authors:** Marina A. Silveira, Yoani N. Herrera, Nichole L. Beebe, Brett R. Schofield, Michael T. Roberts

**Affiliations:** Kresge Hearing Research Institute, Department of Otolaryngology – Head and Neck Surgery, University of Michigan, Ann Arbor, Michigan; Department of Neuroscience, Development and Regenerative Biology, The University of Texas at San Antonio, San Antonio, Texas; University Hospitals Hearing Research Center at NEOMED, Department of Anatomy and Neurobiology, Northeast Ohio Medical University, Rootstown, OH, USA; Department of Molecular and Integrative Physiology, University of Michigan, Ann Arbor, Michigan

**Author notes:** Authors contributed equally (Marina A. Silveira, Yoani N. Herrera).

## Abstract

Growing evidence suggests that neuropeptide signaling shapes auditory computations. We previously showed that neuropeptide Y (NPY) is expressed in the inferior colliculus (IC) by a population of GABAergic stellate neurons and that NPY regulates the strength of local excitatory circuits in the IC. NPY neurons were initially characterized using the NPY-hrGFP reporter mouse, in which hrGFP expression indicates NPY expression at the time of assay, i.e., an expression-tracking approach. However, studies in other brain regions have shown that NPY expression can vary based on a range of factors, suggesting that the NPY-hrGFP mouse might miss NPY neurons not expressing NPY proximal to the experiment date. Here, we hypothesized that neurons with the ability to express NPY represent a larger population of IC GABAergic neurons than previously reported. To test this hypothesis, we used a lineage-tracing approach to irreversibly tag neurons that expressed NPY at any point prior to the experiment date. We then compared the physiological and anatomical features of neurons labeled with this lineage-tracing approach to our prior data set, revealing a larger population of NPY neurons than previously found. In addition, we used optogenetics to test the local connectivity of NPY neurons and found that NPY neurons routinely provide inhibitory synaptic input to other neurons in the ipsilateral IC. Together, our data expand the definition of NPY neurons in the IC, suggest that NPY expression might be dynamically regulated in the IC, and provide functional evidence that NPY neurons form local inhibitory circuits in the IC.

## Introduction

Neuropeptides are small molecules expressed by a large number of neurons throughout the brain and are often co-released with classical fast neurotransmitters such as GABA and glutamate (van den Pol, 2012; Urban-Ciecko and Barth, 2016; Guillaumin and Burdakov, 2021). The co-release of fast neurotransmitters and neuropeptides allows a prolonged reconfiguration of neuronal circuits since neuropeptides exhibit a slower modulatory role by activating metabotropic receptors (Colmers and Bleakman, 1994; Sun et al., 2003; Fu et al., 2004; Nusbaum et al., 2017). In addition, neuropeptides can act over longer distances from the release site, thereby shaping neuronal circuits in ways that are not possible with fast neurotransmitters alone (Nässel, 2009; van den Pol, 2012).

Neuropeptide Y (NPY) is a 36 amino-acid peptide that has been implicated in the modulation of feeding behavior, fear, anxiety, seizures, pain perception, stress responses, and memory (Woldbye et al., 1996; Britton et al., 1997; Gutman et al., 2008; Gøtzsche and Woldbye, 2016; Pedroso et al., 2016; Reichmann and Holzer, 2016; Li et al., 2017; Boyle et al., 2023). NPY has also been used as a molecular marker to identify neuron classes in several brain regions (Karagiannis et al., 2009; Polgár et al., 2011; Silveira et al., 2020). However, because NPY expression can vary over time, there is a distinction between a cell’s ability to express NPY and a cell’s expression of NPY at the time experiments are performed. This raises interesting questions regarding whether the state of a cell should be considered when classifying a cell as an NPY neuron. For example, in the dorsal medial hypothalamus and in the arcuate nuclei, the expression of NPY changes with energy status and food restriction (Brady et al., 1990; Li et al., 1998; Wu et al., 2014; Pedroso et al., 2016). In the auditory brainstem, noise exposure increases the proportion of lateral olivocochlear (LOC) neurons that express NPY (Frank et al., 2023). Additionally, NPY expression can change across development (Botchkina et al., 1996; Morara et al., 1997; Grove et al., 2001; Boyle et al., 2023).

Although the role of NPY in auditory processing remains unknown, NPY is expressed in at least two regions in the subcortical auditory pathway: the LOC (Frank et al., 2023) and the inferior colliculus (IC)(Silveira et al., 2020). The IC forms the major part of the auditory midbrain and is essential for many auditory computations including sound localization and speech processing (Litovsky et al., 2002; Carney et al., 2015; Keine et al., 2016). In the IC, NPY neurons were identified using the NPY-hrGFP mouse line, in which hrGFP expression reports NPY expression at the time of the experiment. Using this expression-tracking approach, our group showed that NPY-expressing neurons represent a distinct class of stellate GABAergic neurons, accounting for nearly one third of IC inhibitory neurons (Silveira et al., 2020). However, because expression of NPY is dynamic and can be influenced by extrinsic factors, it is likely that an expression-tracking approach underestimated the number of neurons that should be counted as NPY neurons.

Here, we hypothesized that the NPY neuron class represents a larger population of IC GABAergic neurons than previously reported and that this expanded population remains homogeneous in its physiological and anatomical properties. To test this hypothesis, we compared two methods for labeling NPY neurons: a lineage-tracing approach and an expression-tracking approach. First, we used NPY-FlpO x Ai65F mice to irreversibly tag neurons that expressed NPY at any point prior to the experiment day, including during development. We refer here to neurons labeled through this lineage-tracing approach as NPY^flp^ neurons. We then compared the properties of NPY^flp^ neurons to our previous dataset obtained from the NPY-hrGFP mouse line in which hrGFP expression reflects active expression of NPY. We refer to these latter neurons, which are labeled through an expression-tracking approach, as NPY^gfp^ neurons. Our data reveal that NPY^flp^ neurons represent a larger population of IC neurons than NPY^gfp^ neurons, due to the inclusion of neurons that were not expressing NPY at the time of experiments. Despite the larger number of NPY^flp^ neurons, both labeling approaches identified neurons with similar physiological and morphological properties, suggesting that they are part of the same neuron class. Since most neurons in the IC have local axon collaterals (Oliver et al., 1991), we next hypothesized that NPY neurons send local inhibitory projections to other IC neurons. Using optogenetic circuit mapping, we found that NPY^flp^ neurons provide inhibitory inputs to other neurons in the local IC. Together, our data indicate that the NPY neuron class in the IC should be expanded to include a set of GABAergic neurons not actively expressing NPY and that NPY neurons form local inhibitory circuits in the IC. These results suggest that NPY expression in the IC is dynamically regulated by intrinsic or extrinsic factors or by developmental states.

## Materials and Methods

### Animals

All experiments were approved by the University of Michigan Institutional Animal Care and Use Committee and were in accordance with the National Institutes of Health’s *Guide for the Care and Use of Laboratory Animals*. Mice were kept on a 12 h day/night cycle with ad libitum access to food and water. To visualize NPY neurons, we used either NPY-hrGFP mice (Jackson Laboratory, stock #006417) (van den Pol et al., 2009) in which NPY^gfp^ neurons are identified by the expression of hrGFP or NPY-FlpO x Ai65F mice in which NPY^flp^ neurons are identified by tdTomato expression by crossing Npy-IRES2-FlpO-D mice (Jackson Laboratory, stock #030211)(Daigle et al., 2018) with Ai65F mice (Jackson Laboratory, stock #032864)(Daigle et al., 2018). All mice were on a C57BL/6J background, and mice of both sexes were used for all experiments.

### Comparison of the distribution of NPY neurons

To evaluate the distribution and density of fluorescent cells, ten NPY-hrGFP mice and four NPY-FlpO x Ai65F mice were deeply anesthetized in an isoflurane drop jar and then perfused transcardially with phosphate-buffered saline (PBS) for 30 – 60 s followed by 10% neutral buffered formalin (Sigma Millipore, catalog #HT501128) for 10-15 min until ∼100 ml of formalin was perfused. Brains were collected and postfixed for 2h, then stored in PBS with 20% sucrose until sectioning. Brains were sectioned on a freezing microtome into 40 µm thick sections. One series (every third section) from each brain was mounted onto gelatin-coated slides, air dried, and coverslipped with DPX mountant (Sigma 317616).

Photomicrographs were obtained with a Zeiss AxioImager.Z2 fluorescence microscope and Hamamatsu Orca Flash 4.0 camera using Zeiss Zen Pro software. Low magnification photomicrographs were collected with a 2.5x objective lens. High magnification photomicrographs were collected as z-stacks using a 63x oil-immersion objective lens (NA = 1.4) with optical sections collected at 0.26 µm depth intervals. Structured illumination with an Apotome 3 (Zeiss) was used to provide optical sectioning. Maximum intensity projections were created from each z-stack.

In nine NPY-hrGFP mice and three NPY-FlpO x Ai65F mice, fluorescent cells were plotted through the rostro-caudal extent of the IC using Neurolucida Software (MBF Biosciences) and a Zeiss AxioImager.Z2 fluorescence microscope with a Hamamatsu Orca Flash 4.0 camera. We plotted 24,609 cells across 9 NPY-hrGFP mice and 33,911 cells across 3 NPY-FlpO x Ai65F mice. IC subdivisions were differentiated with separate tissue series stained for GAD67 and GlyT2 proteins (see Silveira et al., 2020 for details). Plots, counts of cells, and areas for each subdivision were exported using Neurolucida Explorer (MBF Biosciences). Plots were prepared with Adobe Illustrator and numerical analyses were performed and bar graphs were created in Microsoft Excel.

### Fluorescent *in situ* hybridization

The NPY-FlpO x Ai65F mouse line was validated using an RNAscope assay for fluorescent *in situ* hybridization with probes targeted to *tdTomato, Npy* and *Vgat* (*Slc32a1*; Advanced Cell Diagnostics, catalog # 317041, 313321, and 319191)(Wang et al., 2012). Brains were prepared using the fresh frozen method. In brief, two female mice (P51) and one male mouse (P51) were deeply anesthetized using isoflurane and then rapidly decapitated. Brains were quickly harvested, immediately frozen on dry ice, and kept at -80 °C until the day of slicing. After equilibrating the brains at -20 °C, 15 μm coronal sections were collected on a cryostat at −20 °C and mounted on Superfrost Plus slices (Fisher Scientific, catalog # 22037246). Slices were fixed using 10% neutral-buffered formalin (Sigma-Aldrich, catalog # HT501128) and dehydrated in increasing concentrations of ethanol. To block endogenous peroxidase activity, slides were incubated in hydrogen peroxide for 10 min at room temperature followed by application of Protease IV for 30 min. For hybridization, probes targeted to *tdTomato*, *Npy* and *Vgat,* as well as positive and negative controls, were incubated for 2 hours at 40 °C. After the amplification of the probes, the signal was developed using the HRP appropriate for each channel. Opal dyes (1:1000) were assigned for each channel: *Vgat* expression was identified by Opal 520 (Akoya Bioscience, catalog # FP1487001KT), *Npy* expression was identified by Opal 570 (Akoya Bioscience, catalog # FP1488001KT) and *tdTomato* expression was identified by Opal 690 (Akoya Bioscience, catalog # FP1497001KT). Slices were counterstained with DAPI and coverslipped using ProLong Gold antifade mountant (Fisher Scientific, catalog # P36934). Images were acquired within two weeks of completing the assay. Representative sections (caudal, middle and rostral, 4 slices per mouse) were imaged at 2 µm depth intervals with a 0.75 NA 20X objective on a Leica TCS SP8 laser scanning confocal microscope (Leica Microsystems). To quantify the percentage of co-labeling between cells expressing *tdTomato*, *Npy* and *Vgat*, images were exported to Neurolucida 360 (MBF Bioscience), and quantification was performed manually by placing a marker on the top of each cell expressing the mRNA of interest. Cells in each channel were quantified separately to avoid bias in the analysis. DAPI staining was used to verify whether the mRNA puncta were labeling a cell body.

### Immunohistochemistry and analysis

One male P45 and one female P65 NPY-FlpO x Ai65F mouse were deeply anesthetized in an isoflurane drop jar and then perfused transcardially with PBS for 30 – 60 s followed by 10% neutral buffered formalin (Sigma Millipore, catalog #HT501128) for 10-15 min until ∼100 ml of formalin was perfused. Brains were collected and postfixed for 2h. Brains were cut into 40 µm sections on a vibrating microtome (VT1200S, Leica Biosystems). Immunofluorescence using an anti-GAD67 antibody was performed as previously described (Beebe et al., 2016; Goyer et al., 2019; Silveira et al., 2020). In brief, after being washed in PBS, sections were treated with 10% normal donkey serum (Jackson ImmunoResearch Laboratories, catalog #017-000-121) plus 0.3% Triton X-100 for 2 h. Sections were then incubated in a mouse anti-GAD67 antibody (1:1000; Sigma Millipore, catalog #MAB5406, RRID:AB_2278725) for ∼40h at 4°C. On the following day, sections were rinsed in PBS and incubated in Alexa Fluor-647-tagged goat anti-mouse IgG (1:100; Thermo Fisher Scientific, catalog #A21235, RRID:AB_2535804) for 1.5h at room temperature. Brain sections were mounted on Superfrost Plus microscope slides (Thermo Fisher Scientific, catalog #12-550-15) and coverslipped using Fluoromount-G (SouthernBiotech, catalog #0100–01). Images were collected using a Leica TCS SP8 laser scanning confocal microscope with a 1.30 NA 40x oil-immersion objective.

Four coronal IC sections were quantitatively analyzed: two caudal, one mid rostral-caudal and one rostral. Images of one side of the IC were collected at 2 µm Z intervals. Analysis was performed using Neurolucida 360 (MBF Bioscience) by marking the top of each counted cell. Since the anti-GAD67 antibody does not penetrate the entire depth of the tissue sections (Beebe et al., 2016; Goyer et al., 2019), analysis was restricted to regions in which antibody staining was clear, usually the top 10-12 µm of the surface for each section. Fluorescent channels (tdTomato and Alexa 647) were analyzed separately to limit bias.

### Brain slice preparation

To characterize the intrinsic physiology of NPY neurons we performed whole-cell patch-clamp electrophysiology experiments in brain slices from male and female mice aged P35-P80. Recordings were targeted to NPY neurons identified in NPY-FlpO x Ai65F (NPY^flp^) or NPY-hrGFP (NPY^gfp^) mice by the expression of tdTomato or hrGFP, respectively. Data from the NPY-hrGFP mice were previously shown in Silveira et al., 2020. Before decapitation, mice were deeply anesthetized with isoflurane. Dissection of the IC was performed in 34°C ACSF containing the following (in mM): 125 NaCl, 12.5 glucose, 25 NaHCO_3_, 3 KCl, 1.25 NaH_2_PO_4_, 1.5 CaCl_2_, 1 MgSO_4_, 3 sodium pyruvate, and 0.40 L-ascorbic acid, bubbled to a pH of 7.4 with 5% CO_2_ in 95% O_2_. Coronal brain sections containing the IC (200 µm) were prepared using a vibrating microtome (VT1200S, Leica Biosystems). Slices were incubated at 34°C for 30 min in ACSF bubbled with 5% CO_2_ in 95% O_2_ before being transferred to the recording chamber.

### Electrophysiological recordings

Brain slices were maintained at 34°C and continuously perfused at ∼2 ml/min with ACSF bubbled with 5% CO_2_ in 95% O_2_. Neurons expressing either hrGFP or tdTomato were identified with epifluorescence using a Nikon FN1 microscope or an Olympus BX51WI microscope. Recordings were performed in current-clamp mode using BVC-700A patch-clamp amplifiers (Dagan Corporation). Data were low-pass filtered at 10 kHz and sampled at 50 kHz with a National Instruments PCIe-6343 data acquisition board. Data acquisition was done using custom software written in IgorPro (Wavemetrics).

Recording pipettes were pulled in a P-1000 microelectrode puller (Sutter Instrument) using borosilicate glass pipettes (outer diameter 1.5 mm, inner diameter 0.86 mm, catalog #BF150-86-10, Sutter Instrument). The internal pipette solution contained the following (in mM): 115 K-gluconate, 7.73 KCl, 0.5 EGTA, 10 HEPES, 10 Na_2_-phosphocreatine, 4 MgATP, 0.3 NaGTP, supplemented with 0.1% biocytin (w/v), pH adjusted to 7.3 with KOH and osmolality to 290 mmol/kg with sucrose. Pipette resistance ranged from 2.5 – 5.5 MΩ when filled with the internal solution. To assess input resistance, we delivered a series of 100 ms current steps that hyperpolarized the membrane potential to ∼ -100 mV. Peak and steady-state voltage changes were measured, and the peak (R_pk_) and steady-state (R_ss_) input resistances were calculated. To calculate membrane time constant, we applied 40 – 50, 100 – 300 ms current steps that hyperpolarized the membrane potential by 2-6 mV. The time constant was obtained by fitting a single exponential function to each response and calculating the median time constant. All membrane potential values were corrected for the liquid junction potential (-11 mV). Pipette capacitance and series resistance were compensated using the bridge balance of the Dagan amplifier. Recordings with series resistance above 20 MΩ were excluded from the analysis.

### Post hoc reconstructions of morphology and analysis

Recorded neurons were filled with biocytin (Thermo Fisher Scientific, catalog # B1592) via the recording pipette during whole-cell recordings. After the recording, the pipette was slowly removed to allow the cell membrane to reseal. Brain slices were fixed overnight with 10% formalin and were moved to PBS the next day. Slices were stored at 4°C for up to 4 weeks, then were stained using biocytin-streptavidin histochemistry. Slices were washed in PBS 3 times for 10 minutes each, cell membranes were permeabilized using Triton X-100 in PBS for 2 h, washed again 3 x 10 min in PBS, then stained for 24 h with streptavidin-Alexa Fluor-647 (1:1000, Thermo Fisher Scientific, catalog # S21374) at 4°C. The following day, slices were again washed 3 x 10 min with PBS, fixed in 10% formalin for 1 h, and then washed 3 x 10 min with PBS. Slices were then mounted on slides and coverslipped using Fluoromount-G. *z*-stack tile scan images of streptavidin-Alexa Fluor-647 stained neurons were collected using a 1.4 NA 63x oil-immersion objective on a Leica TCS SP8 laser scanning confocal microscope.

Images of recorded neurons were imported into Neurolucida 360 software (MBF Bioscience) where 3D reconstruction of cell somata and dendritic arbors was performed. The major axes of reconstructed neurons were then measured using procedures we previously developed (Goyer et al., 2019; Silveira et al., 2020). In brief, neuron reconstructions were first rotated and/or flipped so that the dorsal-ventral axis was vertical, with the dorsal direction at the top, and the lateral-medial axis was horizontal, with the lateral direction to the left. To assess whether NPY^flp^ neurons have a stellate morphology, a 3D principal component analysis (PCA) was then performed on the *x, y, z* coordinate set for each neuron. The first and second principal directions from the 3D PCA were used to determine the length and the width of each NPY^flp^ neuron in three dimensions, and a length-to-width ratio was calculated. To assess the orientation and extent of neurons relative to the plane of the isofrequency laminae, a 2D PCA was then performed using the *x* and *y* coordinate set for each neuron. The orientation of the first principal component from the 2D PCA was used to determine the long axis angle of each neuron. The *x* and *y* coordinate sets were also used to measure how far the dendrites of each reconstructed neuron extended perpendicular to a 45° laminar plane.

### Intracranial virus injections

To investigate if NPY^flp^ neurons synapse onto other neurons in the ipsilateral IC, 12 NPY-FlpO mice (5 males and 7 females, aged P25 – P82) were injected in the IC with a recombinant adeno-associated virus (rAAV). We tested four different AAVs and did not find any difference between them, therefore the results are reported together. Mice were injected with one of the four viruses: rAAV1/EF1α1.1-FLPX-rc [Chronos-GFP] (Addgene plasmid #122102, AAV prepared by University of Michigan Vector Core, titer 8.47 x 10^13^ VG/ml (Klapoetke et al., 2014)), rAAV8/nEF-Coff/Fon-ChR2(ET/TC)-EYFP (Addgene #137141-AAV8, titer 2.5 x 10^13^ GC/mL (Fenno et al., 2020)), rAAV5/hSyn Coff/Fon hChR2(H134R)-EYFP (University of North Carolina Vector Core #8476, titer 5.8 x 10^12^ GC/mL (Fenno et al., 2014)) and rAAV8/nEF-ChRmine-oScarlet (Addgene #137160-AAV8, titer 2.0 x 10^13^ GC/mL (Fenno et al., 2020)).

Intracranial injections were performed as previously described (Goyer et al., 2019; Silveira et al., 2020, 2023). After being deeply anesthetized with 3% isoflurane, mice were placed in a stereotaxic base with a homeothermic heating pad. During the surgery isoflurane was lowered to 1 – 2% for maintenance. The analgesic Carprofen (5 mg/kg, CarproJect, Henry Schein Animal Health) was administered subcutaneously. The skull was exposed using a rostrocaudal incision, and a craniotomy was made over the left or right IC using a micromotor drill (K.1050, Foredom Electric) with a 0.5 mm burr (Fine Science Tools). The coordinates used for injections (measured relative to lambda) were either 900 µm caudal, 1000 µm lateral or 900 µm caudal, 1200 µm lateral. The depth of the injections varied between 1750 – 2250 µm (measured relative to skull surface at lambda). Injection pipettes were prepared from glass capillaries (catalog # 3-000-203-G/X, Drummond Scientific) pulled with a P-1000 microelectrode puller (Sutter Instrument). The injection tip was front filled with the virus of interest. The scalp was sutured using Ethilon 6-0 (0.7 metric) nylon sutures (Ethicon) or closed using Vetbond Tissue Adhesive (3M). Lidocaine hydrochloride ointment (2%, 0.5ml, Akorn) was then applied to the wound. Experiments were performed 2 - 5 weeks after injections to allow full expression of the opsins.

### Channelrhodopsin-assisted circuit mapping

After preparation of brain slices, recordings were performed under red room lighting to limit the activation of opsins by ambient light. Chronos or Channelrhodopsin were activated by brief pulses of 470 nm light emitted by a blue LED. ChRmine was activated by brief pulses of green light produced by a white LED filtered with a 545 nm bandpass filter with a ± 12.5 nm pass band (cat # ET545/25x, Chroma Technology). Both LEDs were coupled to the epifluorescence path of the microscope and light was delivered using a 0.80 NA 40x water immersion objective. Optical power densities ranged from 0.9 to 48 mW/mm². Light duration ranged from 1 – 3 ms using a 5 – 30 s inter-sweep interval. Only sweeps that exhibited a light-evoked IPSP were included in the analysis. In a subset of experiments, we tested for pharmacological block of IPSPs by applying 5 µM gabazine (also called SR95531 hydrobromide, GABA_A_ receptor antagonist, Hello Bio, Cat #: HB0901) and 1 µM strychnine hydrochloride (glycine receptor antagonist, Sigma-Aldrich, Cat #: S8753). Drugs were bath applied for at least 5 – 10 minutes before responses were measured.

### Statistics

Statistical analyses were performed using R 4.1.0 (The R Project for Statistical Computing), Igor Pro 9 (Wavemetrics), and MATLAB R2021a (MathWorks). For the comparison of the intrinsic physiology between NPY^gfp^ and NPY^flp^ neurons, we used Welch’s t-test, and the significance level (α) was adjusted to account for multiple comparisons using Bonferroni correction. For the remaining experiments, effects were considered significant when *p* < 0.05. Data are shown as mean ± SD. Principal component analysis (PCA) was performed using the *“coeff*” function in MATLAB. The first component of the PCA explained 97.18% of the data. After PCA, a k-means clustering analysis was performed using the *“kmeans”* function in MATLAB. The “elbow method” was used to determine how many clusters to divide the data into.

## Results

### Most IC neurons labeled in NPY-FlpO x Ai65F mice express *Npy* mRNA and are GABAergic

To test whether the NPY-FlpO x Ai65F mouse line selectively labels IC neurons that express NPY and are GABAergic, we performed *in situ* hybridization using an RNAscope assay with probes targeted to mRNAs encoding *tdTomato, Npy* and the vesicular GABA transporter, *Vgat*. Two females and one male NPY-FlpO x Ai65F mouse, all aged P51, were used for the assay. We found that 75.6% (3111 out of 4115) of *tdTomato^+^* neurons expressed *Npy*, and 92.9% (3822 out of 4115) of *tdTomato^+^* neurons expressed *Vgat* (**Figure 1A-E**, **Table 1**). To confirm the neurotransmitter content of NPY^flp^ neurons, we also performed immunofluorescence against GAD67, an enzyme essential for GABA synthesis and an established marker of GABAergic IC neurons (Ono et al., 2005; Ito et al., 2009; Beebe et al., 2016; Goyer et al., 2019; Silveira et al., 2020). Using representative coronal IC sections from one male (P45) and one female (P65), we found that 92.2% (2333 out of 2528) of tdTomato^+^ neurons labeled with anti-GAD67 (**Figure 1F-H**). The NPY-FlpO x Ai65F mouse line therefore predominately labels a population of GABAergic neurons, three-quarters of which expressed *Npy* at the time of the assay.

**Figure 1.**
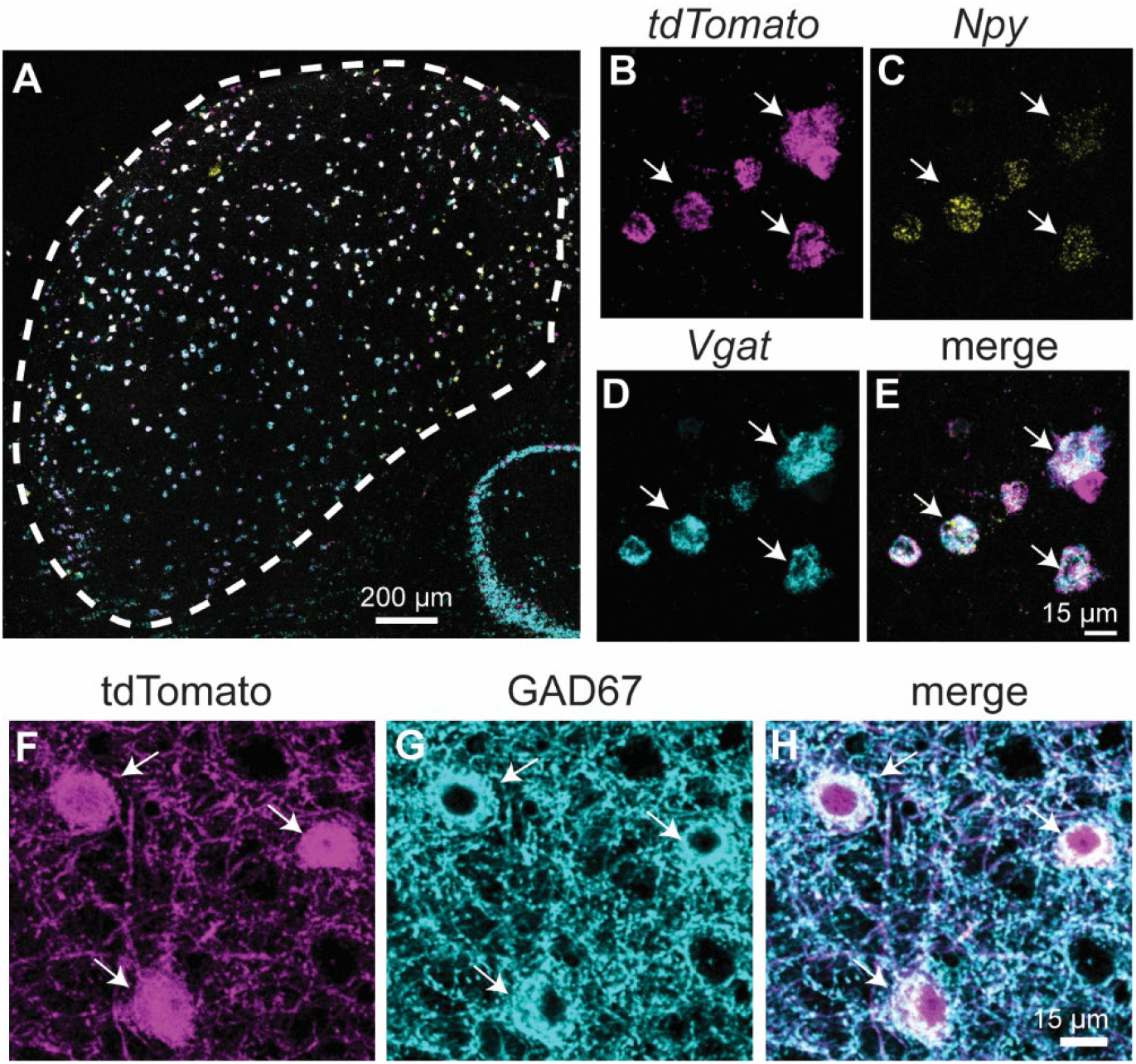
Most *tdTomato^+^*neurons in NPY-FlpO x Ai65F mice express *Npy* and are GABAergic. **A-E.** Fluorescent *in situ* hybridization (RNAscope) was used to determine the expression patterns of *tdTomato*, *Npy* and *Vgat* mRNA in the IC of NPY-FlpO x Ai65F mice. **A.** Low magnification coronal IC section from an NPY-FlpO x Ai65F mouse shows the distribution of *tdTomato^+^* (magenta), *Npy^+^* (cyan) and *Vgat^+^* (yellow) neurons in the IC. **B-E.** High magnification confocal images show that *tdTomato^+^* neurons (**B**, magenta) typically co-labeled with *Npy* (**C**, cyan) and *Vgat* probes (**D,** yellow). Merged image shown in **E**. White arrows highlight examples of neurons that co-labeled. Scale bar in **E** applies to images in **B-E**. **F-H.** Immunofluorescence shows that 92.3% of tdTomato^+^ neurons are GABAergic. High magnification confocal images show tdTomato^+^ neurons (**F**, magenta) immunolabeled with antibodies against the GABA synthetic enzyme, GAD67 (**G**, cyan). Merge in **H**. Scale bar in **H** applies to images in **F-H**.

**Table 1.** Nearly all *tdTomato*^+^ neurons expressed *Vgat* and three-quarters expressed *Npy*.

| Mouse | *tdTomato*<sup>+</sup> | *tdTomato*<sup>+</sup><br>*Npy*<sup>+</sup> | % *tdTomato*<sup>+</sup><br>*Npy*<sup>+</sup>/<br>*tdTomato*<sup>+</sup> | *tdTomato*<sup>+</sup><br>*Vgat*<sup>+</sup> | % *tdTomato*<sup>+</sup><br>*Vgat*<sup>+</sup>/<br>*tdTomato*<sup>+</sup> | *Npy*<sup>+</sup> | % *tdTomato*<sup>+</sup><br>*Npy*<sup>+</sup>/<br>*Npy*<sup>+</sup> |
| --- | --- | --- | --- | --- | --- | --- | --- |
| Female<br>P51 | 154 | 108 | 70.1 | 127 | 82.5 | 117 | 92.3 |
|  | 609 | 464 | 76.2 | 580 | 95.2 | 482 | 96.3 |
|  | 420 | 290 | 69.0 | 414 | 98.6 | 298 | 97.3 |
|  | 271 | 205 | 75.6 | 257 | 94.8 | 230 | 89.1 |
| Female<br>P51 | 498 | 389 | 78.1 | 464 | 93.2 | 398 | 97.7 |
|  | 394 | 317 | 80.5 | 364 | 92.4 | 326 | 97.2 |
|  | 338 | 290 | 85.8 | 305 | 90.2 | 299 | 97.0 |
|  | 465 | 340 | 73.1 | 413 | 88.8 | 348 | 97.7 |
| Male<br>P51 | 345 | 280 | 81.2 | 329 | 95.4 | 298 | 94.0 |
|  | 157 | 93 | 59.2 | 133 | 84.7 | 123 | 75.6 |
|  | 132 | 80 | 60.6 | 127 | 96.2 | 117 | 68.4 |
|  | 332 | 255 | 76.8 | 309 | 93.1 | 266 | 95.9 |
| **Total** | **4115** | **3111** | **75.60** | **3822** | **92.9** | **3302** | **94.22** |

To test whether the NPY-FlpO x Ai65F mouse line reliably labels the population of neurons expressing *Npy* at the time of assay (as opposed to a subset), we next analyzed the *in situ* hybridization data for co-labeling of *Npy^+^* cells. The results showed that 94.2% of *Npy^+^*cells expressed *tdTomato* and 99.4% expressed *Vgat* (**Table 2**). Thus, the NPY-FlpO x Ai65F mouse line labels nearly all *Npy*-expressing cells, and, as we found in our previous study (Silveira et al., 2020), *Npy*-expressing cells are GABAergic.

**Table 2.** Over half of *Vgat*^+^ neurons expressed *tdTomato* and/or *Np*.

| Mouse | *Vgat*<sup>+</sup> | *Vgat*<sup>+</sup><br>*Npy*<sup>+</sup> | % *Vgat*<sup>+</sup><br>*Npy*<sup>+</sup>/<br>*Vgat*<sup>+</sup> | *Vgat*<sup>+</sup><br>*tdTomato*<sup>+</sup> | % *Vgat*<sup>+</sup><br>*tdTomato*<sup>+</sup>/<br>*Vgat*<sup>+</sup> | *Npy*<sup>+</sup> | % *Vgat*<sup>+</sup><br>*Npy*<sup>+</sup>/<br>*Npy*<sup>+</sup> |
| --- | --- | --- | --- | --- | --- | --- | --- |
| Female<br>P51 | 321 | 117 | 36.4 | 127 | 39.6 | 117 | 100.0 |
|  | 822 | 482 | 58.6 | 580 | 70.6 | 482 | 100.0 |
|  | 707 | 297 | 42.0 | 414 | 58.6 | 298 | 99.7 |
|  | 451 | 228 | 50.6 | 257 | 57.0 | 230 | 99.1 |
| Female<br>P51 | 594 | 398 | 67.0 | 464 | 78.1 | 398 | 100.0 |
|  | 499 | 326 | 65.3 | 364 | 72.9 | 326 | 100.0 |
|  | 473 | 298 | 63.0 | 305 | 64.5 | 299 | 99.7 |
|  | 500 | 346 | 69.2 | 413 | 82.6 | 348 | 99.4 |
| Male<br>P51 | 493 | 297 | 60.2 | 329 | 66.7 | 298 | 99.7 |
|  | 322 | 115 | 35.7 | 133 | 41.3 | 123 | 93.5 |
|  | 442 | 115 | 26.0 | 127 | 28.7 | 117 | 98.3 |
|  | 409 | 262 | 64.1 | 309 | 75.6 | 266 | 98.5 |
| **Total** | **6033** | **3281** | **54.38** | **3822** | **63.35** | **3302** | **99.36** |

**Table 3.** Nearly all tdTomato^+^ neurons co-labeled with GAD67.

| Mouse | tdTomato<sup>+</sup> | tdTomato<sup>+</sup><br>GAD67<sup>+</sup> | tdTomato<sup>+</sup><br>GAD67<sup>-</sup> | % tdTomato<sup>+</sup><br>GAD67<sup>+</sup>/<br>tdTomato<sup>+</sup> |
| --- | --- | --- | --- | --- |
| Male P45 | 456 | 423 | 8 | 92.8 |
|  | 745 | 661 | 49 | 88.7 |
| Female<br>P65 | 380 | 337 | 32 | 88.7 |
|  | 947 | 912 | 14 | 96.3 |
| **Total** | **2528** | **2333** | **103** | **92.3** |

Together, these results indicate that the NPY-FlpO x Ai65F mouse line drove tdTomato expression in three populations of neurons under our experimental conditions: 75.1% were GABAergic neurons that expressed *Npy* at the time of assay, 17.8% were GABAergic neurons that did not express *Npy* at the time of assay, and 7.1% were non-GABAergic (presumptive glutamatergic) neurons that did not express *Npy* at the time of the assay (calculations based on the assumption that 99.4% of *tdTomato^+^ Npy^+^* neurons were GABAergic). This differs from our previous study using NPY-hrGFP mice, where 94.7% of NPY^gfp^ neurons expressed NPY (Silveira et al., 2020), and suggests that the lineage-tracking approach of the NPY-FlpO x Ai65F mouse line results in broader labeling than the expression-tracking approach of the NPY-hrGFP mouse line. The broader labeling in the NPY-FlpO x Ai65F mouse line could be due to ectopic expression of tdTomato in non-NPY-expressing neurons, labeling of neurons that expressed NPY at some point prior to the date of assay but not on the assay day, or some combination of the two. Since NPY can be expressed transiently during development and can be regulated by extrinsic factors (Botchkina et al., 1996; Morara et al., 1997), we hypothesize that most of the additional labeling in the NPY-FlpO x Ai65F mouse line was due to labeling neurons that fit the NPY class definition (i.e., GABAergic neurons with sustained firing patterns and stellate morphology) but were not actively expressing *Npy* at the time of assay.

### NPY^flp^ and NPY^gfp^ neurons have similar distributions but different densities in the IC

To analyze the distribution and density of labeled cells in each mouse line, we examined fluorescently labeled NPY cells in the IC of both NPY-hrGFP and NPY-FlpO x Ai65F mice. **Figure 2A** shows low-magnification images of the IC in NPY-hrGFP (green, left) or NPY-FlpO x Ai65F (magenta, right) mice. In both types of mice, fluorescently labeled cells are readily visible throughout the IC, including in each major IC subdivision and through the caudo-rostral extent of the nucleus. Commissural fibers are fluorescently labeled in each type of mouse (arrows), indicating the participation of NPY cells in the IC commissural pathway. In a major difference between the two types of mice, GABA modules of the IClc (Chernock et al., 2004) are visible in the NPY-FlpO x Ai65F mouse IC (arrowheads) but not visible in the NPY-hrGFP mouse IC. A second difference between the two types of mice is the increased density of fluorescent cells in the NPY-FlpO x Ai65F mouse IC. This increased density was also visible at higher magnification (**Figure 2B**). The density of fluorescent cells was increased in the NPY-FlpO x Ai65F mouse IC as compared to the NPY-hrGFP mouse IC in each major subdivision of the IC. This was especially visible in the IClc (**Figure 2B**, bottom row), where clusters of NPY^flp^ cells were present within GABA modules. As at lower magnifications, IC commissural fibers are readily visible in the ICd of each type of mouse (**Figure 2B**, middle row), although fibers appeared brighter (compared to somatic fluorescence) and denser in the NPY-hrGFP mouse IC.

**Figure 2.**
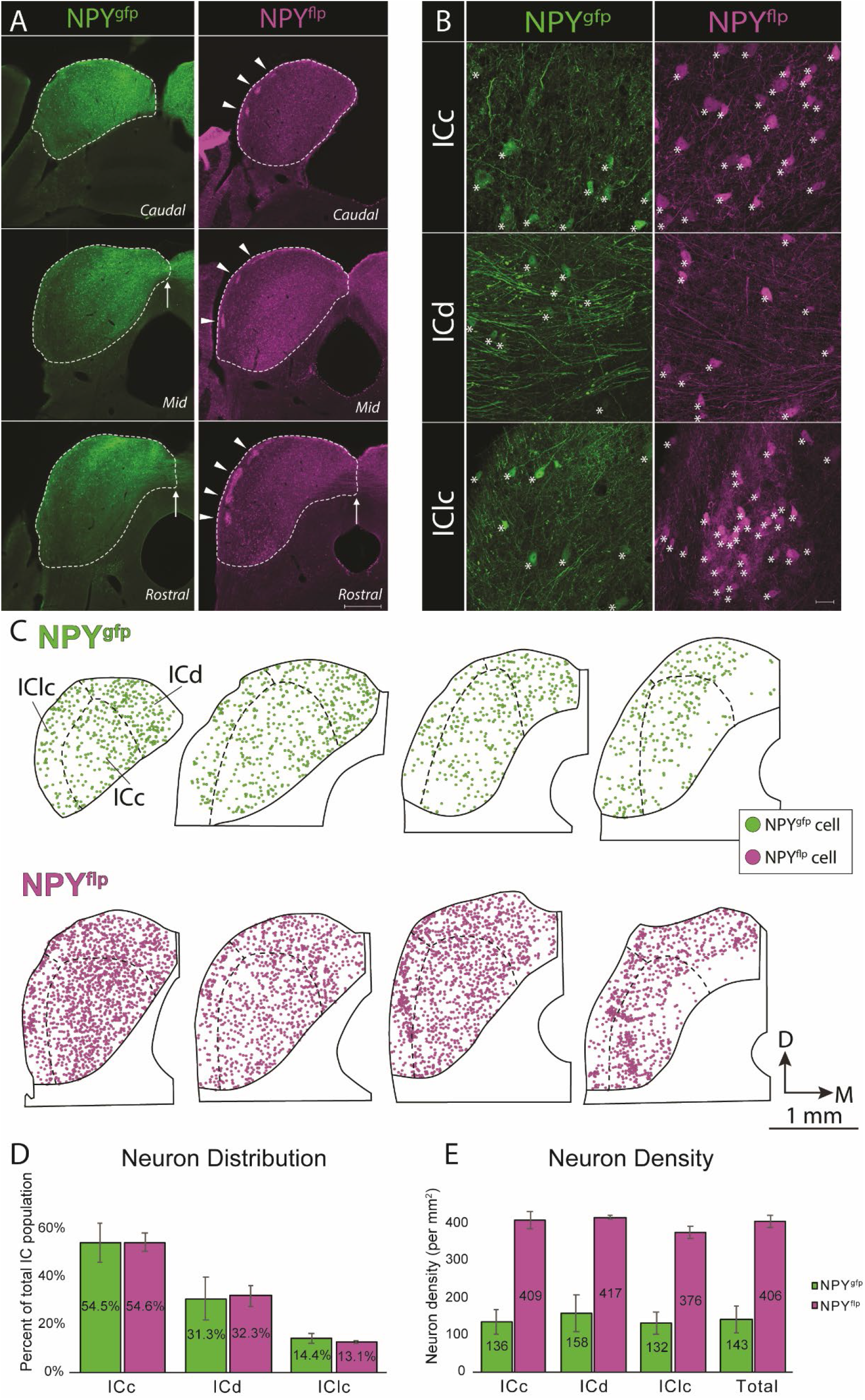
NPY^flp^ and NPY^gfp^ neurons have similar distributions but different densities in the IC. **A.** Low-magnification photomicrographs showing NPY^gfp^ (left column, green) or NPY^flp^ (right column, magenta) fluorescence in the IC. For each type of mouse, sections from the caudal (top row), mid (middle row) or rostral (bottom row) IC are shown. Note that commissural fibers can be seen in both types of mice (arrows), while GABA modules from the IClc (arrowheads) are only visible in NPY-FlpO x Ai65F mice. Scale = 500 µm. **B.** High-magnification photomicrographs showing NPY^gfp^ (left column, green) or NPY^flp^ (right column magenta) fluorescence in each IC subdivision. Fluorescent cells (denoted with asterisks) are readily visible in each subdivision in both types of mice (top row: ICc, middle row: ICd, bottom row: IClc). However, cells were more numerous in a given field in each subdivision in NPY-FlpO x Ai65F mice. Note especially the presence of many NPY^flp^ cells in a GABA module in the IClc. Fluorescent axons and boutons are also readily visible in each subdivision in both types of mice, and commissural axons are especially prominent in the ICd of both types of mice. Scale = 20 µm. **C.** Plots showing each fluorescent cell in four representative sections in one case from each type of mouse. One green circle denotes one NPY^gfp^ cell and one magenta circle denotes one NPY^flp^ cell. For both cases, sections are arranged from caudal (left) to rostral (right). D - dorsal; M - medial. **D-E.** Bar graphs showing the percentage of total population in each subdivision (**D**) and density of cells per mm^2^ in each subdivision and overall (**E**) for NPY^gfp^ cells (green bars) and NPY^flp^ cells (magenta). Bars show mean +/- SD. n = 24,609 NPY^gfp^ cells across 9 NPY-hrGFP mice and 33,911 NPY^flp^ cells across 3 NPY-FlpO x Ai65F mice.

To further investigate the difference in density of fluorescent cells between the two models, we plotted each fluorescently labeled cell through the IC in nine NPY-hrGFP mice and three NPY-FlpO x Ai65F mice. Representative sections from each model are shown in **Figure 2C**, where each circle represents one fluorescent cell. We found no difference in the relative distribution of labeled cells between the IC subdivisions when we compared the percentage of the total population present in each subdivision (**Figure 2D**). However, when we compared the density of fluorescent cells (cells per mm^2^ of subdivision area), we found a much higher density of NPY^flp^ as compared to NPY^gfp^ neurons in the IC (**Figure 2E**). This combination of higher density and similar distribution seems to indicate no major differences in the cell populations that are fluorescently labeled in the two types of mice, however subtle differences do seem to exist (for example, clusters of NPY^flp^ cells in IClc modules that are not labeled in the NPY-hrGFP mouse IC).

### The intrinsic physiology of NPY^flp^ neurons is similar to NPY^gfp^ neurons

Neurons in the IC exhibit heterogenous intrinsic physiological properties (Peruzzi et al., 2000). Previously, we showed that NPY^gfp^ neurons exhibit a sustained firing pattern, low expression of hyperpolarization-activated cation current (*I_h_*) and moderate input resistances and membrane time constants (Silveira et al., 2020). Because our *in situ* hybridization results suggested that the NPY-FlpO x Ai65F mouse labels a larger population of neurons than the NPY-hrGFP mouse line, we hypothesized that the intrinsic physiology of this larger population remains homogeneous compared to the NPY^gfp^ neurons. To test this hypothesis, we performed whole-cell current clamp recordings targeted to NPY^flp^ neurons and compared their intrinsic physiology with our previous data for NPY^gfp^ neurons. The firing pattern of each neuron was classified based on the spike frequency adaptation ratio (SFA) in response to a depolarizing current step, such that neurons with an SFA > 2 were classified as adapting and neurons with an SFA < 2 were classified as sustained (Peruzzi et al., 2000). We found that 85% of NPY^flp^ neurons exhibited a sustained firing pattern (n = 50/59), similar to the 95% we saw previously for NPY^gfp^ neurons (n = 123/129; **Figure 3A,B**). The mean SFA ratios for NPY^flp^ neurons trended higher than those of NPY^gfp^ neurons (NPY^flp^ neurons= 1.73 ± 0.78; NPY^gfp^ neurons = 1.26 ± 0.66; mean ± SD), but this difference was not statistically significant (SFA ratio for current steps eliciting 5 spikes comparing NPY^flp^ *vs* NPY^gfp^ neurons: Welch’s t-test, *p =* 0.056. Since six physiological parameters were compared for the same cells for the experiment shown in **Figure 3**, the Bonferroni-corrected α = 0.008).

**Figure 3.**
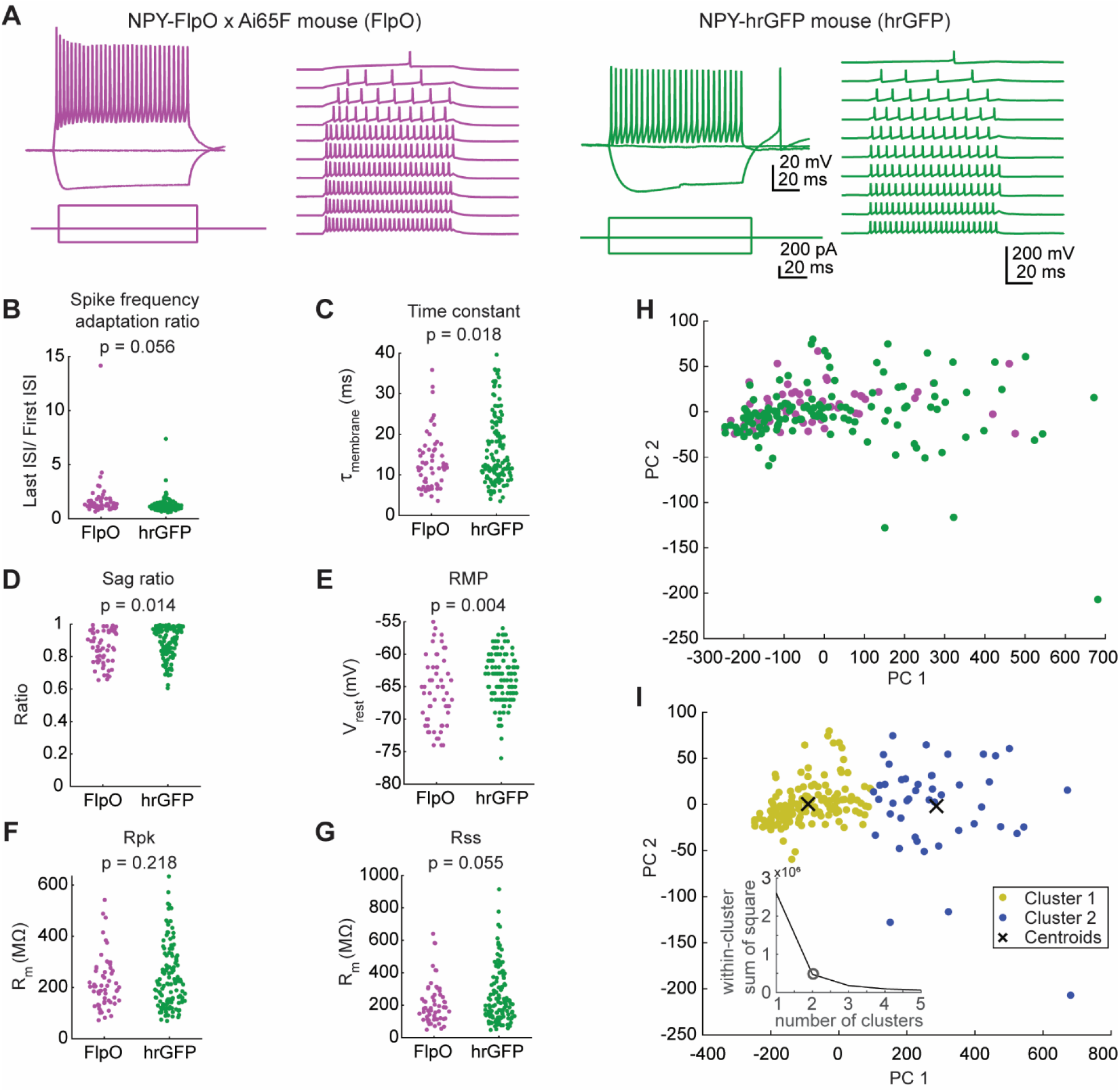
NPY^flp^ and NPY^gfp^ neurons exhibit similar intrinsic physiological properties. **A.** NPY^flp^ (magenta) and NPY^gfp^ (green) neurons exhibited sustained firing patterns. **B-G**. NPY^flp^ and NPY^gfp^ neurons exhibited similar spike frequency adaptation ratios (**B**), membrane time constants (**C**), voltage-dependent sag ratios (**D**), resting membrane potentials (**E**), input resistances measured from the peaks of a series of hyperpolarizing responses (**F**), and input resistances measured from the steady state portions of a series of hyperpolarizing responses (**G**). *p* values from Welch’s t-tests are shown atop each plot. The right side of each graph shows a paired mean difference plot (Gardner-Altman estimation plot), with the horizontal black line indicating the paired mean difference (hrGFP minus FlpO). The vertical black line indicates the 95% confidence interval for the paired mean difference as calculated from a bootstrap analysis, with the gray distribution showing the distribution of mean differences obtained from the bootstrap analysis (see Methods for details). Dashed gray lines represent the level of zero difference. Final statistical determinations were based on the Welch’s t-test, but paired mean difference plots are provided to give a better sense of the distribution of differences between the pairs of experimental groups. **H**. Principal component analysis showed that the distributions of NPY^flp^ (magenta) and NPY^gfp^ (green) neurons largely overlapped. **I.** k-means cluster analysis identified two clusters of neurons from the PCA results, represented by yellow and blue dots (dot positions are identical to **H**). Both clusters contained NPY^flp^ and NPY^gfp^ neurons, with the yellow cluster containing 86.4% of NPY^flp^ neurons and 71.3% of NPY^gfp^ neurons, showing that NPY^flp^ and NPY^gfp^ neurons were not separable based on their intrinsic physiology. The number of clusters used for analysis was defined using the elbow analysis method, which showed that the addition of a third cluster did little to improve the separation between cluster centroids (inset graph).

To calculate membrane time constant, negative current steps were used to hyperpolarize the membrane potential 2 – 6 mV, and 40 – 50 sweeps were recorded. An exponential function was fit to each sweep and the median τ was obtained. The membrane time constant of NPY^flp^ neurons was similar to that of NPY^gfp^ neurons (13.1 ± 6.9 ms vs 15.9 ± 8.5 ms; Welch’s t-test, *p* = 0.018. Bonferroni-corrected α = 0.008, **Figure 3C**). When injected with a hyperpolarizing current step, NPY^flp^ and NPY^gfp^ neurons exhibited moderate sag ratios suggesting relatively low expression of *I_h_* (0.85 ± 0.10 vs 0.89 ± 0.09; Welch’s t-test, *p* = 0.014; Bonferroni-corrected α = 0.0083, **Figure 3D**). The resting membrane potential was slightly more hyperpolarized for NPY^flp^ neurons compared to NPY^gfp^ neurons (-65.8 ± 5.1 mV vs -63.5 ± 3.8 mV; Welch’s t-test, *p* = 0.004; Bonferroni-corrected α = 0.008), but the effect size was small (mean difference of 2.8 mV, **Figure 3E**). Finally, input resistance measured at the peak (R_pk_) or at the steady state of the hyperpolarizing response (R_ss_) was similar between NPY^flp^ and NPY^gfp^ neurons (R_pk_: 216.9 ± 100.8 MΩ vs 237.8 ± 120.1 MΩ; Welch’s t-test, *p* = 0.055; Bonferroni-corrected α = 0.008, **Figure 3F**. R_ss_: 209.6 ± 129.0 MΩ vs 252.7 ± 167.6 MΩ; Welch’s t-test, *p* = 0.2186; Bonferroni-corrected α = 0.008, Figure 3G).

Next, we performed PCA analysis to test whether NPY^flp^ and NPY^gfp^ neurons have overlapping or separable distributions based on intrinsic physiology. PCA analysis was conducting by combining the NPY^flp^ and NPY^gfp^ data sets for the six intrinsic physiology parameters assessed above: SFA ratio, membrane time constant, voltage sag ratio, resting membrane potential, R_pk_, and R_ss_. The results from the PCA analysis suggested that NPY^flp^ and NPY^gfp^ neurons have largely overlapping distributions (**Figure 3H**). We next performed k-means cluster analysis that was set to identify 2 clusters (the optimal number of clusters defined by the “elbow method,” see insert in **Figure 3I**). We found that 86.4% of NPY^flp^ neurons and 71.3% of NPY^gfp^ neurons were part of cluster 1 while the remaining neurons fell into cluster 2, suggesting that the two populations of NPY neurons were similarly divided between the two clusters (**Figure 3I**). These data suggest that the intrinsic physiology of NPY^flp^ and NPY^gfp^ neurons is similar, supporting the hypothesis that both neuron groups originate from the same class of neurons.

### NPY^flp^ neurons have stellate morphology similar to NPY^gfp^ neurons

We reconstructed the morphology of 15 recorded NPY^flp^ neurons using biocytin-streptavidin staining. **Figure 4A** shows all reconstructed neurons, all of which were located in the ICc. For comparison purposes, all neurons were oriented as they would appear in the left side of a coronal IC slice viewed from the caudal side.

**Figure 4.**
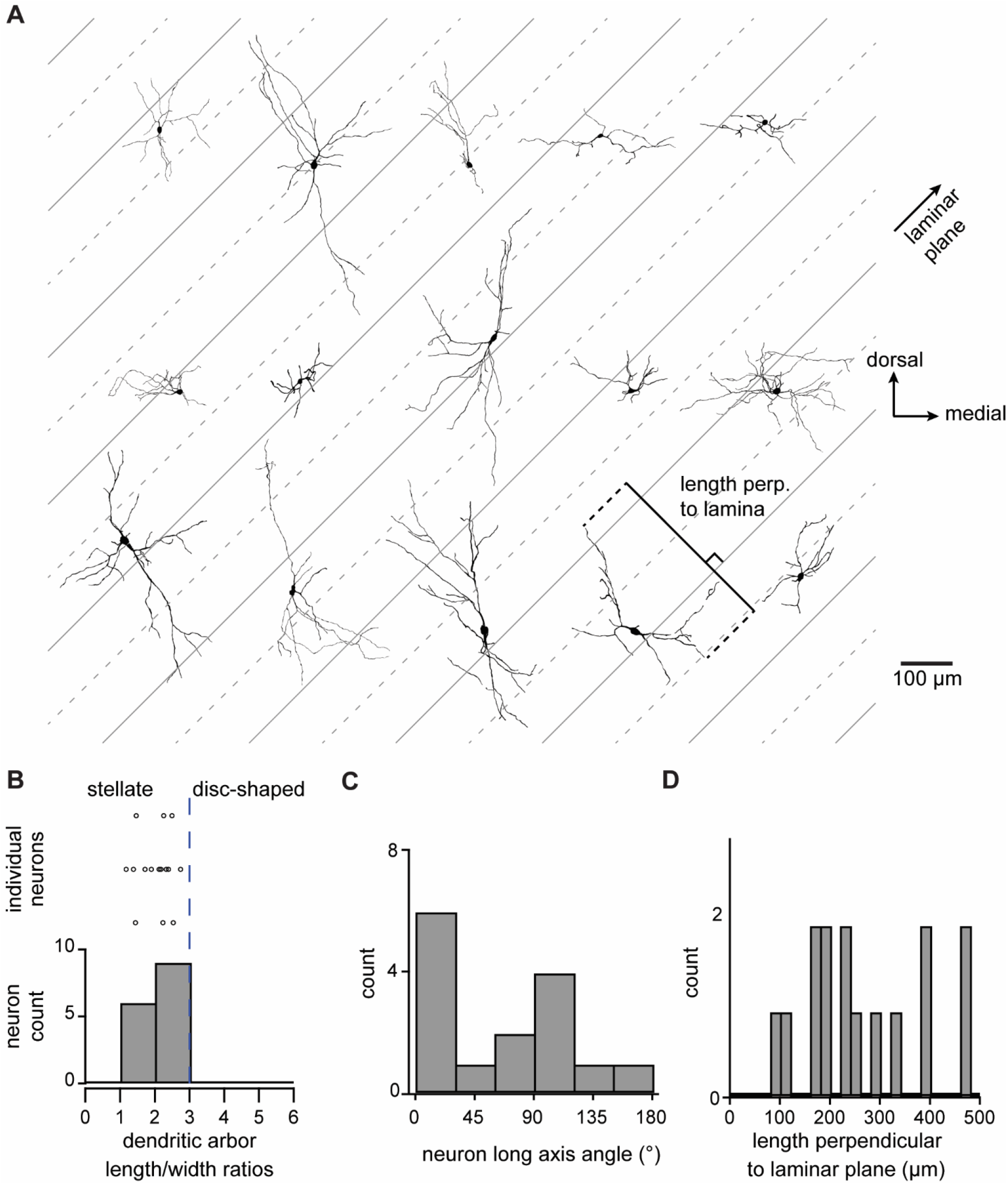
NPY^flp^ neurons in the ICc have stellate morphology similar to NPY^gfp^ neurons. **A.** Representative reconstructions of the morphology of 15 NPY^flp^ neurons that were filled with biocytin during recordings from the ICc. Gray lines represent the approximate orientation and width of ICc isofrequency lamina (∼45° angle and 100 µm, respectively). **B.** 3D PCA analysis was performed to classify neuron morphology using the ratio of the longest and second-longest axes of each neuron (i.e., length and width), with a ratio <3 indicating stellate morphology. **C.** Angular orientation of the long axis of reconstructed neurons. Angles indicate counter-clockwise rotation of the long axis relative to the medial-lateral (horizontal) axis of the IC such that a 45° angle is parallel to the isofrequency plane. **D.** Length dendritic arbors of NPY^flp^ neurons extended perpendicular to the isofrequency laminae plane.

First, using an approach developed in our earlier studies (Goyer et al., 2019; Silveira et al., 2020), we used 3D PCA to classify the morphology of each neuron as stellate or disc-shaped. This analysis was done on the x, y, z coordinate set for each NPY^flp^ neuron to determine the extent along the first principal direction (length) and extent along the second principal direction (width) of each neuron in three dimensions. We then calculated the length-to-width ratio of the dendritic arbor of each neuron.

Previous work has shown that stellate neurons have a length-to-width ratio < 3 and disc-shaped neurons have a ratio of ≥ 3 (Oliver et al., 1991). We found that all reconstructed neurons had a length-to-width ratio < 3, consistent with the hypothesis that NPY^flp^ neurons are stellate shaped (**Figure 4B**).

In the mouse ICc, isofrequency laminae extend at a ∼45° angle relative to the horizontal plane, and disc-shaped neurons are expected to have a long axis that extends parallel to the isofrequency laminae (Morest and Oliver, 1984; Stiebler and Ehret, 1985). As in our previous studies (Goyer et al., 2019; Silveira et al., 2020), we used 2D PCA to calculate the orientation of the long axis of NPY^flp^ neuron dendritic arbors in the coronal plane in relation to ICc isofrequency laminae. The results show that reconstructed NPY^flp^ neurons did not have a preferred orientation, and only one neuron had an orientation that was parallel to isofrequency laminae (**Figure 4C**).

Because the dendrites of ICc neurons typically extend across more than one isofrequency lamina, we measured the distance ICc NPY^flp^ neurons extended their dendrites perpendicular to a 45° laminar plane. We found that only 1 of 15 NPY^flp^ neurons had a dendritic arbor that extended < 90 µm perpendicular to the laminar plane and only 2 of 15 had dendritic arbors that extended < 150 µm perpendicular to the laminar plane (**Figure 4D**). This suggests that most NPY^flp^ neurons have dendrites that extend across two or more isofrequency lamina, with some branching across more than five laminae.

### NPY^flp^ neurons form local inhibitory circuits in the IC

Most neurons in the IC are thought to project to extrinsic targets and to also have local axon collaterals, allowing them to contribute to local circuits (Oliver et al., 1991; Ito et al., 2016). We recently showed that neurons that express the NPY Y_1_ receptor form interconnected excitatory networks in the local IC (Silveira et al., 2023). However, whether NPY neurons contribute to local IC circuits is still unknown. Because most techniques for manipulating distinct neuron types are optimized to target neurons that express Cre and/or FlpO transgenes (Fenno et al., 2011, 2014, 2020), the NPY-hrGFP mouse line is not amenable for selectively manipulating NPY neurons. By contrast, to test if NPY^flp^ neurons form functional connections within local IC circuits, we were able to use FlpO-dependent adeno-associates viruses (AAVs) to selectively express excitatory channelrhodopsins in NPY-FlpO mice (see Methods for details).

Although NPY^flp^ neurons are distributed throughout the IC, our data showed that most neurons expressing excitatory opsins (Chronos-GFP, ChR2-EYFP, ChR2-mCherry or ChRmine-mScarlet) following Flp-dependent AAV transfections were restricted to the edge of the IC (**Figure 5A**). This result suggests that despite strong expression of NPY in the IC, the FlpO expression in the IC is likely to be low in this mouse line. To test whether channelrhodopsins drive firing in NPY neurons, we first targeted recordings to cells expressing the fluorophore. We found that presentation of light elicited action potential trains (**Figure 5B).** We next targeted recordings to other neurons near the transfected area and found that 1-5 ms pulses of blue light elicited inhibitory post synaptic potentials (IPSPs) in 13 out of 28 cells recorded (peak amplitude = -2.0 ± 1.3 mV, mean ± SD, **Figure 5C,D**). In a subset of neurons (n = 6) we subsequently applied gabazine (5 µM) and strychnine (1 µM) to block GABA_A_ and glycine receptors. In all cases the light evoked IPSP was abolished in the presence of these drugs (amplitude in control condition: -1.8 mV ± 0.9 mV; amplitude in gabazine + strychnine: 0 mV; Welch’s t-test, *p* = 0.0005, **Figure 5E,F**). These data indicate that NPY^flp^ neurons contribute to local circuits in the IC by providing functional inhibitory input to other IC neurons.

**Figure 5.**
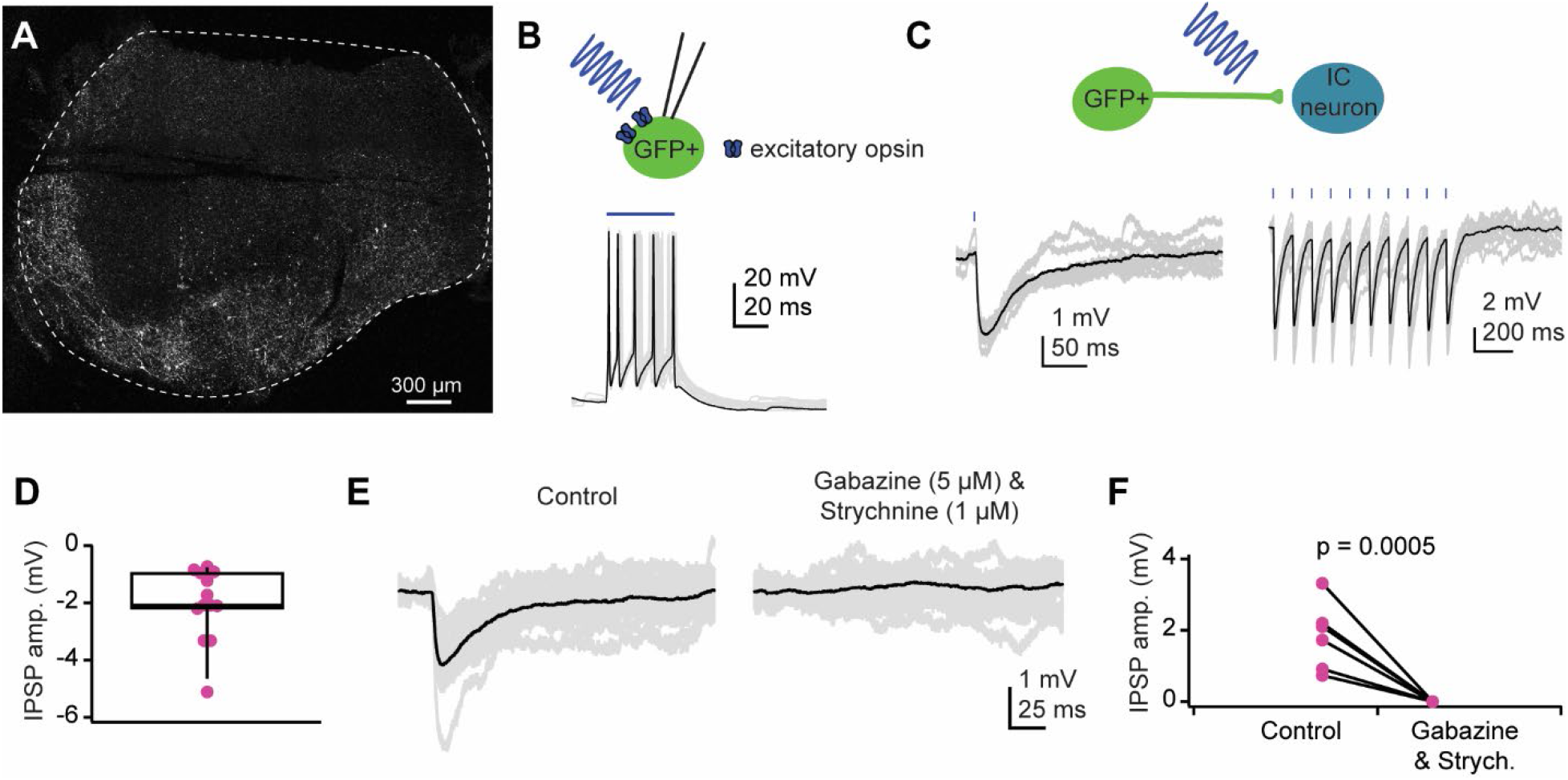
NPY^flp^ neurons provide inhibitory input to IC neurons in the ipsilateral IC. **A.** Confocal image showing the typical expression pattern observed when using a FlpO-dependent channelrhodopsin virus to transfect IC neurons in an NPY-FlpO mouse. This example shows transfection with AAV1/EF1α1.1-FLPX-rc [Chronos-GFP]. **B.** Representative example of a whole-cell recording targeted to a GFP^+^ cell from a mouse injected in the IC with a FlpO-dependent Chronos-GFP virus (AAV1/EF1α1.1-FLPX-rc [Chronos-GFP]). Presentation of a blue light pulse elicited action potentials, confirming the ability to use optogenetics to drive NPY neuron firing. Black trace highlights a representative trial, and gray traces show additional individual trials. **C.** Representative example of optogenetically evoked IPSPs elicited by activating NPY^flp^ neurons and/or terminals in the local IC. Brief flashes of blue light could be used to elicit single IPSPs or trains of IPSPs (20Hz light presentation shown). **D.** Distribution of IPSP amplitudes observed across all recorded cells that received input from NPY^flp^ neurons. **E.** Representative example trace of optogenetically evoked IPSPs recorded from a neuron that received input from an NPY^flp^ neuron (left). The response was abolished with application of GABA_A_ and glycine receptor antagonists (right). **F.** Application of gabazine (5 µM) and strychnine (1 µM) abolished light-evoked IPSPs in all cells. In **C,E,** gray traces represent individual trials and black traces represent averages across trials.

## Discussion

Here we compared lineage-tracing and expression-tracking approaches for identifying NPY neurons in the IC. We found that the lineage-tracing, NPY-FlpO x Ai65F mouse line labels an expanded set of neurons in the IC compared to the expression-tracking, NPY-hrGFP mouse line. While the percentages of neurons expressing NPY mRNA or peptide differed between mouse lines (75.6% NPY^flp^ vs 94.7% NPY^gfp^), the neurotransmitter content, anatomical distribution, and morphological and physiological features of fluorophore-labeled neurons remained similar between mouse lines. Our data therefore suggest that neurons actively expressing NPY and neurons that previously expressed NPY belong to the same class of neurons. Since NPY expression can be regulated by extrinsic factors, we propose that NPY neurons in the IC comprise both neurons actively expressing NPY and neurons with the potential to express NPY.

In addition, using targeted optogenetics, we found that NPY neurons provide inhibitory input to other neurons in the ipsilateral IC, expanding the repertoire of postsynaptic targets for NPY neurons beyond the commissural and tectothalamic projections previously shown. Thus, we conclude that NPY neurons represent a larger than previously known class of inhibitory IC neurons that contribute to local and long-range inhibitory circuits and in which NPY expression might be dynamically regulated.

### Lineage-tracing reveals a larger population of NPY neurons in the IC

Across brain regions, it is well established that different classes of neurons play different roles in neuronal computations (Petilla Interneuron Nomenclature Group et al., 2008; Pelkey et al., 2017; Zeng and Sanes, 2017; Zeng, 2022). Understanding the organization and function of neuronal circuits therefore requires the ability to identify and manipulate distinct neuron types. Although the IC is a major site of auditory processing, it was not until recently that molecularly identifiable neuron types have been identified in the IC (Beebe et al., 2016; Goyer et al., 2019; Silveira et al., 2020; Kreeger et al., 2021). Among those neuron types, NPY neurons were the first molecularly identifiable class of GABAergic neurons (here called NPY^gfp^ neurons) (Silveira et al., 2020). Here we show that a lineage-tracing approach reveals a larger population of neurons that fit the NPY class definition than the expression-tracking approach used in our previous study. These results could be interpreted in two ways. First, they could indicate that NPY is transiently expressed during development in a subset of IC neurons and that these neurons are not able to express NPY in adulthood. If this interpretation is correct, NPY^flp^ neurons would likely be divided into two different classes of neurons: one consistent with the NPY^gfp^ neurons previously described and another representing a previously unknown neuron type.

However, our data challenge this hypothesis as NPY^flp^ neurons had anatomical, morphological, and physiological features similar to NPY^gfp^ neurons. While our morphological analyses showed that all the NPY^flp^ neurons we reconstructed had stellate morphology, a few neurons had distinctly smaller dendrites than others. This could be a result of the dendritic arbors of these neurons being cut when sliced for experiments, but it could also indicate heterogeneity within the NPY neuron population. Consistent with this, the Allen Brain Cell Atlas identifies 44 clusters of GABAergic neurons in the IC, 15 of which express NPY (Yao et al., 2023).

Second, differences in the number of labeled NPY neurons could suggest that NPY expression in the IC is dynamic and influenced by extrinsic factors. Dynamic changes in NPY expression have been described in other brain regions. For example, in the hypothalamus, NPY expression changes with food restriction (Brady et al., 1990; Li et al., 1998; Pedroso et al., 2016) and, in the hippocampus and frontal cortex, NPY release is enhanced during seizures (Marksteiner and Sperk, 1988; Marksteiner et al., 1989; Gruber et al., 1994; Noe’ et al., 2007). In the auditory brainstem, the number of lateral olivocochlear efferent neurons expressing NPY increased after noise exposure (Frank et al., 2023). Similarly, NPY expression in the IC might increase during and/or following periods of heightened activity, which might be caused by noisy environments, audiogenic seizures, and other stimuli. In addition, the IC exhibits enhanced central gain after hearing loss (Chambers et al., 2016; Shaheen and Liberman, 2018), and therefore noise trauma might also induce changes in NPY expression. Since NPY signaling dampens the excitability of most glutamatergic IC neurons (Silveira et al., 2020, 2023), increased NPY expression following hearing loss might help keep enhanced central gain from driving hyperexcitability in the IC.

These findings also raise the importance of carefully planning experiments when using Cre/Flp/Dre recombinase-driver mouse lines. For example, in the dorsal horn, the role of NPY neurons was initially underestimated because viral approaches used to selectively target NPY neurons transfected neurons that transiently expressed NPY during development as well as neurons that expressed NPY in adulthood (Bourane et al., 2015). It was not until recently, using a viral strategy to transfect only neurons that expressed NPY during adulthood, that it was recognized that dorsal horn NPY neurons have a broader role in pain and itch (Boyle et al., 2023). Similarly, for future studies using the NPY-FlpO mouse line to label NPY neurons in the IC, it will be important to consider the developmental stage when Flp-dependent virus transfections are performed and possibly also the environmental conditions in which mice are housed (e.g., ambient noise levels).

Finally, we note that 7.1% of tdTomato-expressing neurons in the NPY-FlpO x Ai65F mouse line were negative for *Vgat* mRNA expression. There are several possible explanations for this. First, there may be some leakage of FlpO expression in the NPY-FlpO mouse such that some non-GABAergic neurons express FlpO. Second, NPY might be transiently expressed during development in a small population of non-GABAergic IC neurons. Third, the anti-*Vgat* probe used in the RNAscope assay might not have labeled all *Vgat*-expressing cells. While these are important considerations to keep in mind, in the realm of recombinase-driver mouse lines, where expression patterns are rarely perfect, the ∼93% efficacy of the NPY-FlpO x Ai65F mouse line in labeling GABAergic IC neurons and our observation that NPY^flp^ and NPY^gfp^ neurons had highly similar properties suggest that the NPY-FlpO mouse is a useful tool for studying IC NPY neurons.

### Considerations for using FlpO-driver mouse lines

Despite the large number of neurons expressing *Npy* in Npy-FlpO x Ai65F mice (∼75% of *tdTomato^+^* neurons expressed *Npy* mRNA), the Flp-dependent viral transfections we conducted produced fluorescence in relatively few neurons, and this fluorescence was more prevalent in the shell regions of the IC. This pattern was observed with all four viruses tested suggesting that it was not dependent on the virus serotype, viral promoter, or the excitatory opsin being expressed (see Methods for list of viruses tested). This result suggests that FlpO expression in P25 - P82 mice (the age range when virus injections were conducted) is stronger in IC shell regions than in the ICc in the NPY-FlpO mouse line. This expression pattern could be useful for future studies focused on local inhibitory circuits in the shell IC. However, we cannot rule out the possibility that NPY^flp^ neurons in the ICc also expressed excitatory opsins but at lower levels than were detectable on our electrophysiology rig microscope.

On the other hand, a recent study provided evidence that FlpO has poorer recombinase activity than Cre, which may explain the relatively poor expression of Flp-dependent viral constructs we observed in NPY-FlpO mice (Zhao et al., 2023). While this may be a disadvantage for experiments aiming to uniformly label NPY neurons using viral transfections, it may create an opportunity for experiments aiming to tie reporter gene expression levels to driver gene expression levels (Zhao et al., 2023). For example, since manipulations that enhance NPY expression should also enhance FlpO expression in NPY-FlpO mice, the expression of Flp-dependent viral constructs might change following noise exposure, audiogenic seizures, hearing loss, or other manipulations that might alter NPY expression. It will be interesting to see whether this feature of FlpO can be leveraged in future studies.

### NPY neurons contribute to local IC circuits

The IC is rich in local circuits and most IC neurons are thought to have local axon collaterals (Oliver et al., 1991; Saldana and Merchan, 2005; Chen et al., 2018). Using glutamate uncaging, previous studies showed that IC GABAergic neurons contribute to local IC circuits, and the distribution of intrinsic inhibitory connections changed with development and noise exposure (Sturm et al., 2014, 2017). We previously showed that at least a portion of NPY neurons are principal neurons that project to the auditory thalamus and/or the contralateral IC (Silveira et al., 2020; Anair et al., 2022). Here, using channelrhodopsin-assisted circuit mapping, we found that NPY neurons can also provide inhibitory synaptic input to other neurons in the local IC. However, it is still unknown whether individual NPY neurons project to multiple targets or if subsets of NPY neurons project to different targets.

In the IC, synaptic inhibition influences frequency tuning and direction selectivity for frequency modulated (FM) sweeps, both of which are important for processing speech and other vocalizations (Kuo and Wu, 2012; Pollak, 2013). In addition, diminished synaptic inhibition in the IC is a common feature in aging and age-related hearing loss (Caspary et al., 1995; Ibrahim and Llano, 2019). Since NPY neurons represent a large proportion of IC GABAergic neurons (Silveira et al., 2020), inhibitory inputs from NPY neurons may be well positioned to shape frequency tuning, FM direction selectivity, and other auditory computations. In addition, NPY neurons might use the co-release of GABA and NPY to regulate local circuits over spatial and temporal scales that are not possible with GABA release alone. For example, we showed that ∼80% of IC glutamatergic neurons express the NPY Y_1_ receptor, and these neurons are hyperpolarized by exogenous application of NPY (Silveira et al., 2023). Therefore, NPY release is expected to dampen the excitability of most glutamatergic neurons in the IC, potentially improving the precision of auditory computations and acting as a brake on hyperexcitability.

Together, our data show that NPY neurons in the IC encompass a larger population of IC GABAergic neurons than previously reported, that NPY expression might be developmentally and/or dynamically regulated in these neurons, and that NPY neurons contribute to local inhibitory circuits in the IC. Future studies will focus on determining what drives NPY expression in IC neurons.

## Acknowledgements

This work was supported by National Institutes of Health Grants R01 DC018284 (MTR), K99 DC019415 (MAS), F31 DC021618 (YNH), T32 DC000011 (YNH), and the University Hospitals Hearing Research Center at NEOMED.

## Author Contributions

MAS and MTR conceived and designed the research. MAS, YNH, NLB and BRS performed experiments. All authors analyzed data and interpreted the results of the experiments. MAS and YNH wrote the initial draft of the manuscript. MAS, YNH and NLB prepared figures. All authors revised, edited and the approved the final version of the manuscript.

